# Inferring evolutionary distances from whole prokaryotic genome alignments, accounting for among-locus substitution rate heterogeneity

**DOI:** 10.64898/2026.10.15.759355

**Authors:** Paul Etheimer, Tommaso Stentella, Anaelle Hulbert, Peter F. Arndt, Chloé-Agathe Azencott, Michael Sheinman, Florian Massip

## Abstract

Ever since the rise of next generation sequencing technologies, the number of prokaryotic genomes in databases has increased tremendously. In order to leverage those vast quantities of data, an approach was previously introduced to estimate evolutionary divergences by modeling the genomic distances between substitutions in many whole genome alignments. However, this proof of concept study was conducted on the *Enterobacterales* clade only, such that its applicability to other clades remained to be established. In this study, we demonstrate the broad applicability of this method to a wider range of clades and to species with larger evolutionary divergences. We also show that relaxing a hypothesis on the substitution rate distribution, with a single additional free parameter, drastically improves both the estimation of evolutionary divergences and the quality of the inferred trees. We use simulations to show that our method can correctly recover the tree topology in cases where substitution rates evolve along the tree. Finally, our results also have theoretical implications for our understanding of substitution rate heterogeneity in genomes, since we found differences in the range of substitution rate distribution in our three examples *Bacillales, Enterobacterales* and *Methanobacteriota* whose causes remain to be identified.

## 1 Introduction

Using molecular sequences to infer evolutionary histories has been an important tool of evolutionary biology for many decades (Zuckerlandl, Emil and Pauling, Linus, 1965). In the genomic age, more and more data are at scientists’ disposal, and progress in sequencing, assembly and annotation of genomes helps researchers uncover uncharted areas of biodiversity. However, data accumulation comes first, and, for some tasks such as phylogenomics, method progress lags behind. Indeed genomes are usually composed of segments of discordant history, because of transfers, duplication, losses or incomplete lineage sorting, which complicates the inference of a phylogeny with standard approaches. Furthermore, some taxa of life like Bacteria or even more so, Archeae, are less well described and understood than model species or mammals. Phylogenetic information is nonetheless essential in order to comprehend, and potentially leverage, the wealth of information contained in the genomes of those less-studied domains of life.

Multiple sequence alignment is key to most approaches in phylogenomics, but its large compute time requirements can be a limitation when studying thousands of genomes, such that most approaches use only one genome per species. But, in the case of Bacteria, leveraging the diversity of the pangenome (Vernikos et al., 2015) could help use more information than a single representative genome. To tackle the challenge posed by increasing quantity of sequence data, alignment-free distance methods have been developed (Criscuolo, 2019; Katz et al., 2019; Zielezinski et al., 2019b; Wang et al., 2024). One drawback of alignment-free methods is their lack of formal modeling of evolutionary processes, such as Horizontal Gene Transfer (HGT) in the case of prokaryotes, that can lead to underestimation of evolutionary distances (Zielezinski et al., 2019a). Other, more heavily parameterized inference approaches have been improved to take into account realistic biological scenarios such as HGT, gene duplication and loss, substitution rate heterogeneity or evolution (Boussau et al., 2013; Morel et al., 2024; Weiner et al., 2025; Zhang et al., 2025). However, those models usually scale poorly with the number of species, and work well only when genome annotation is of good quality. Recently, simulation-based inference (Lueckmann et al., 2021) has enabled the use of more complex models. Because of such complexity, likelihood-based optimization is not a feasible inference approach and researchers resort to either using ABC-based approaches (Torrance et al., 2024; He et al., 2026) or using neural networks (Nesterenko et al., 2025; Min et al., 2026). But these methods require an extremely large amount of simulated data (half a million simulations, requiring hundred of hours of CPU time, for (Torrance et al., 2024)), leading to significant training costs in the case of neural networks.

To address the lack of explicit model of alignment-free method and the computing time and annotation requirements of multiple sequence alignment approaches, we proposed a method (Sheinman et al., 2024), that we here call `mosaic-pair`. `mosaic-pair` co-estimates speciation time and HGT rate between pairs of bacterial species, while still leveraging the increasing amount of information in databases. This method represents pairwise whole genome alignments as Match Length Distributions (MLDs), the distribution of distances between consecutive substitutions, contrasting with other distance methods that measure or estimate the number of mismatches over the alignment length (Konstantinidis and Tiedje, 2005; Jain et al., 2018; Criscuolo, 2019; Shaw and Yu, 2023). It leverages the abundance of genomic data and helps disentangle conflicting signals of genome evolution in bacteria, namely horizontally transferred sequences biasing distance estimation. Entire genomes can be used without the complex task of finding widely shared and transfer-free loci before the species tree inference, unlike what is most commonly done (Kapli et al., 2020) for character-based methods. Indeed, the MLD of a pair of genomes can be used to distinguish the two processes creating matches, clonal inheritance and horizontal transfer. Furthermore, this model allows for among-locus substitution rate heterogeneity (a “mosaic” of molecular clocks) by assuming a genome-wide rate distribution, hence remaining frugal in terms of parameters. Compared to distance methods, this approach models aligner sensitivity, HGT and rate heterogeneity to obtain estimates for both transfer rates and distance between pairs of species.

It was hence previously shown that information on pairwise HGT rates and time divergences is contained in MLDs. As a proof-of-concept, the `mosaic-pair` model was used to infer a tree of *Enterobacterales* Sheinman et al., 2024). However this model does not generalize well to other clades. First, it assumes fixed rate distribution parameters, which do not hold for other species, and second the reasoning behind the shape of the substitution rate distribution conflicts with evolution along a tree. Additionally, we did not measure the performance of `mosaic-pair` at tree reconstruction. In this study, we fitted rate distribution parameters from the collection of pairwise species comparisons, extending the previous approach into a new model, `mosaic-tree`. We also devised a simulation framework in order to characterize the performance and robustness of the tree estimation. Testing the `mosaic-tree` model on several prokaryotic clades, we found that it was able to recapitulate phylogenetic relationships in all tested cases, generalizing beyond *Enterobacterales*. `mosaic-tree` therefore bridges the gap between pure distance methods and character-based methods by modeling the distribution of substitution rates at the tree level while inferring distances between pairs of species.

## 2 Results

### 2.1 Model overview

The `mosaic-pair` and `mosaic-tree` methods represent pairwise alignments of whole genomes as Match Length Distributions (MLDs). Genome evolution is inferred from MLDs, modeled with the continuous stick-breaking process (Arndt, 2019), which was previously used in comparative genomics to study duplication and conservation (Massip and Arndt, 2013; Massip et al., 2015, 2016). The main assumptions of the model are then the following: 1. because substitution rates vary among loci, vertically inherited genomic regions shared by a pair of species *i* do not all have the same evolutionary divergence (defined as the expected number of substitutions per site along the path between the two tips). We assume that these divergences follow an approximately linear distribution, from *θ*_*i*_/*κ* for the most conserved regions to *θ*_*i*_ for regions under no selection. This generalizes the molecular clock to rate variation along genomes (see Supp. Mat. 10.3 and 10.4). 2. the rate of transfer is constant through time and exchanged pieces of DNA have identical substitution rates; 3. substitutions appear as a random Poisson process on infinite number of sites. Justification for these are to be found in (Sheinman et al., 2024) and supplementary. Additionally, in `mosaic-tree` we use dimensionless units (see Supp. Mat. 10.4) to better separate tree inference and time calibration.

Consequently, MLDs observed while comparing bacterial species’ genomes present two regimes arising from two processes. Following assumption 1., the distribution of segment divergences gives rise to a first regime, for short matches, where the frequency of match lengths decreases in proportion to the −4 power of their length. Following assumption 2., a constant transfer rate and a unique substitution rate for transferred segments leads to a uniform distribution of divergences for this part of the genome. It gives rise to a second regime, for longer matches, where the frequency of matches decreases in proportion to a −3 power of their length. For most MLDs, the vertical regime dominates for the shortest matches as most of the inter-species matches arise from vertical inheritance, while longer matches are dominated by recent HGT events, as such long matches are very unlikely to be preserved uninterrupted since speciation.

The `mosaic-pair` model assumed a predetermined substitution rate distribution for the vertical regime, with boundaries fixed from the literature. Indeed we used studies of *Escherichia coli* (Wielgoss et al., 2011) to fix the boundaries of the substitution rate, which was used to report a time *τ*. In contrast, in mosaic-tree *κ* is shared by all pairs and is fitted alongside all the pairwise transfer rates and distances *θ*_*i*_. This allows `mosaic-tree` to be more flexible and adapted to the studied clade. Additionally, the aligner used in `mosaic-pair`, MUMmer (Marçais et al., 2018), was not sensitive enough and was replaced in `mosaic-tree` with lastz (Harris, 2007) (see 4.4 for more details). The `mosaic-tree` model is summarized in Fig. 1.

**Figure 1:**
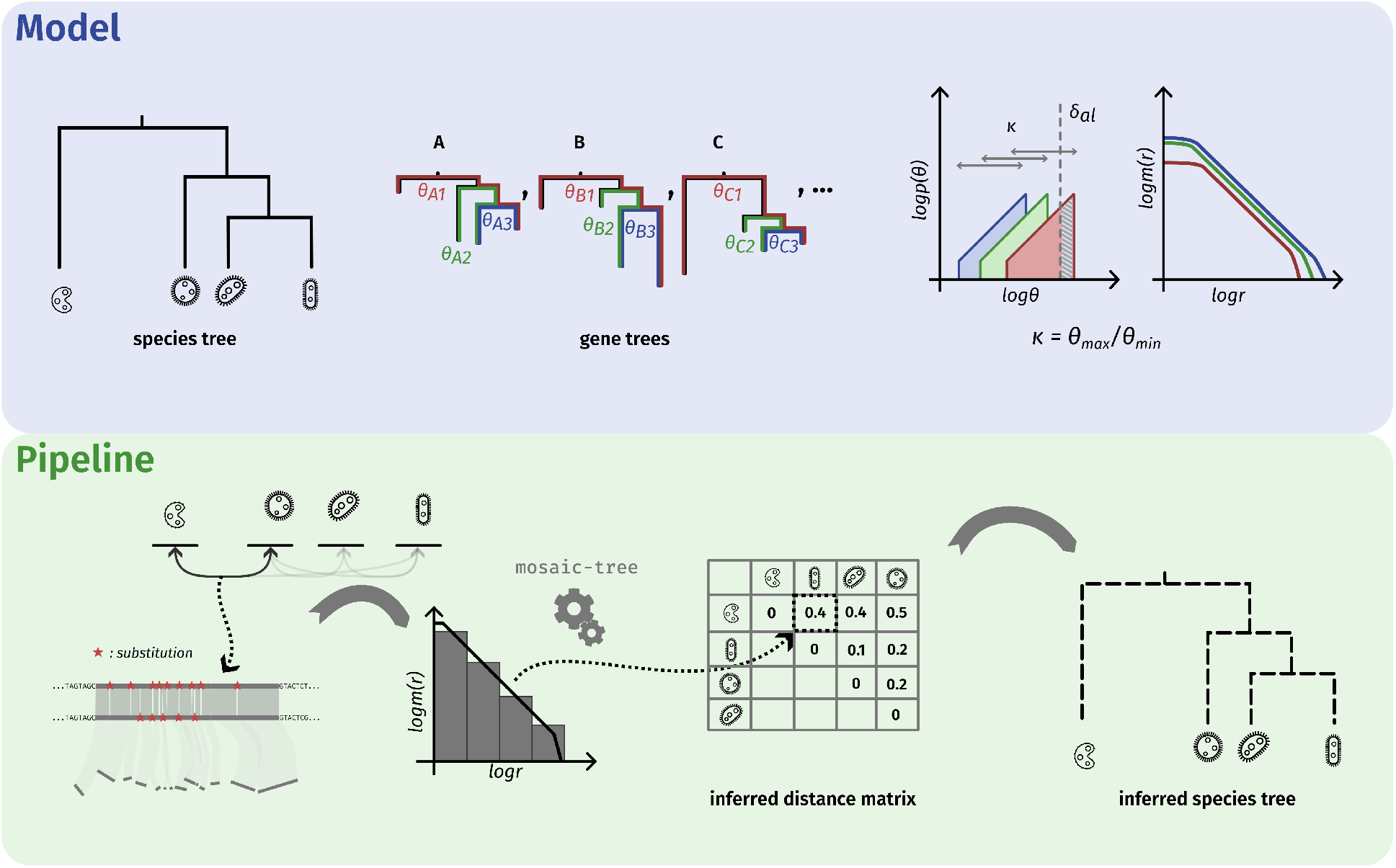
Overview of the mosaic-tree model and pipeline. The model assumes rate heterogeneity in the gene trees (A, B, C… in the figure), leading to a distribution of divergences per species comparison approximated by a linear distribution of width *κ* (top right, left). Those linear distributions give rise to a Match Length Distribution (MLD) following a power law (top right, right). The pipeline starts by pairwise genome alignments, computes the empirical MLDs, infers a matrix of distances, and uses it to build a tree using the UPGMA algorithm (Sokal et al., 1958).

### 2.2 Simulations show that mosaic-tree is able to recover phylogenies accurately

In `mosaic-pair`, we assumed that the linear rate distribution arose from the sum of two uniform rate distributions in the two aligned genomes. For reason developed in Supp. Mat. 10.3, this assumption is not compatible with trees and had to be discarded. On the other hand, doing so unlocked our ability to simulate entire trees.

To determine the ability of the method to infer the additional, tree-wide parameter added to `mosaic-tree`, we simulated MLDs (more details in Material and Methods 4.3) and tested the inference for different amounts of data and tree heights. Rates were drawn from a linear distribution and shared across each gene tree. Results indicate that in most settings, we can estimate *κ* using all pairwise MLDs of the simulated tree (Supp. Fig. S2).

This first simulation scenario is however unlikely to properly describe genome evolution. Apart from essential genes, complete correlation of all genomic region’s rate evolution is unlikely, and a very strong assumption. Indeed, it is equivalent to no discordance between gene trees and species tree for every gene. Additionally, in real settings we do not have access directly to MLDs but to whole genomes. To further evaluate estimation of the parameter *κ* in more realistic settings, we developed a tree-compatible genome evolution simulation pipeline. In this second scenario, substitution rates evolve through a bounded random walk, reproducing the strict bounds of the rate distribution of our model.

Applying `mosaic-tree` to data simulated with this second approach, we obtained trees whose topology is perfectly recovered by the inference. As opposed to our first simulation scenario, we find that distances are consistently underestimated (Figure 2). The systematic underestimation indicates that the linear approximation of the rate distribution is probably incorrect in this case. Despite this shortcoming, `mosaic-tree` was able to recover the ordering of the pairwise distances, thus the topology.

**Figure 2:**
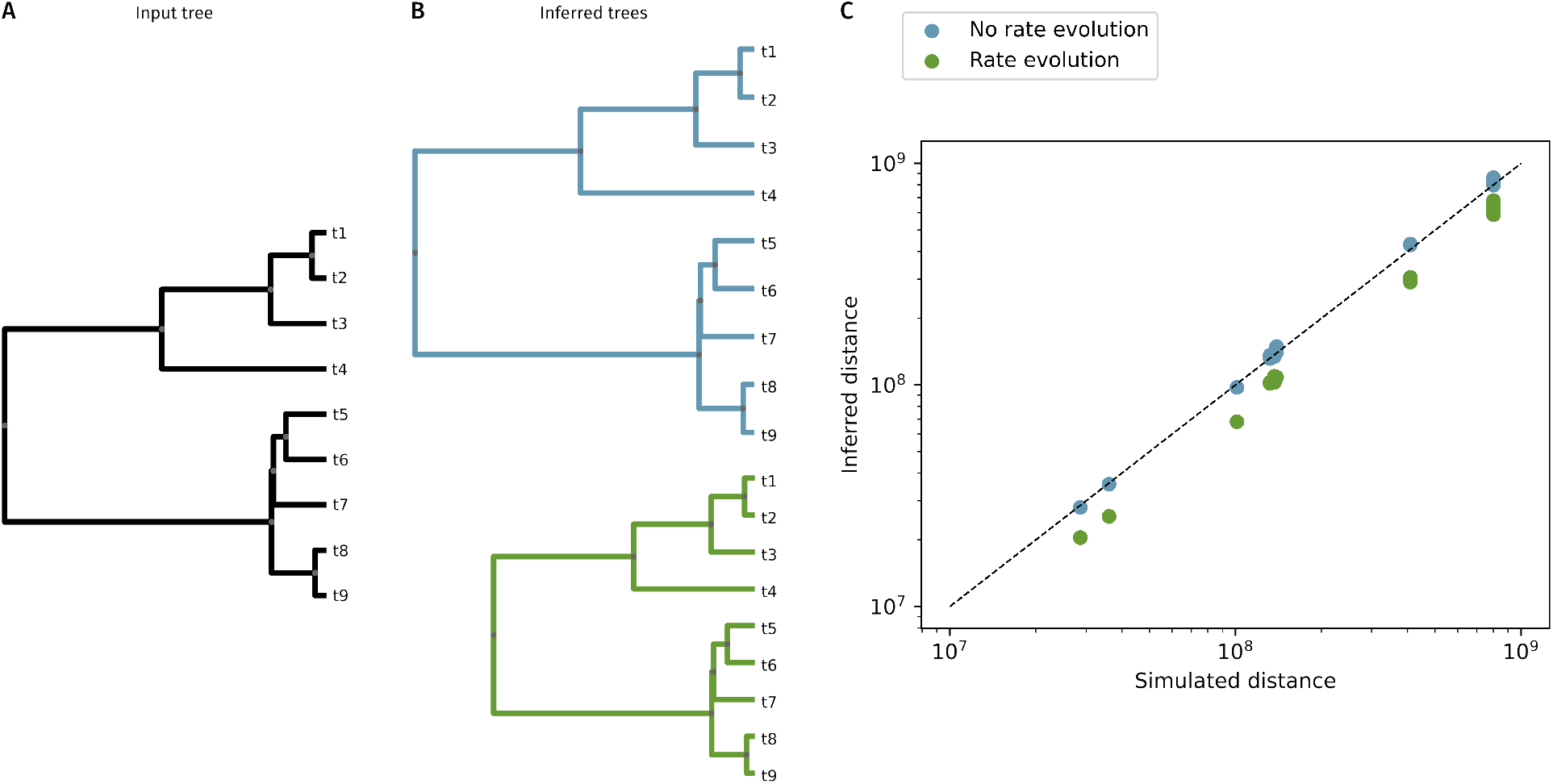
Simulation and inference of a random tree. A: A tree is simulated in two scenarios, with gene substitution rate drawn from a common linear rate distribution and with Brownian substitution rate evolution. B: In the first (blue, top) and second (green, bottom) scenarios, `mosaic-tree` correctly recovers the topology. C: In the first scenario, `mosaic-tree` recovers the distances, while in the second scenario, the distances are consistently underestimated. Trees are drawn to scale.

### 2.3 Fits of mosaic-tree are better

In order to study the wider applicability of the mosaic method, we tested it on other clades in addition to *Enterobacterales*. We selected the bacterial order *Bacillales* (as defined by the Genome Taxonomy Database (GTDB) (Parks et al., 2022), see 4.1), an important clade of the Gram+ model bacteria *Bacillus subtilis* with plenty of genomes in databases. We also studied Archaea, and selected the archeal phylum *Methanobacteriota* as this is the archeal clade with the highest number of genomes available in GTDB.

First, we find that `mosaic-tree` fitted MLDs better than `mosaic-pair`, especially for short matches (Fig. 3 A and B). The least improved fits (Fig. 3 C) were still well fitted with this new model, indicating that using it might be beneficial, or at least never detrimental, for every species comparisons. We note however that in those cases, there could be a tradeoff between fitting a shorter value of *κ* across all comparisons and fitting a higher transfer rate in the cases where this value of *κ* is too small.

**Figure 3:**
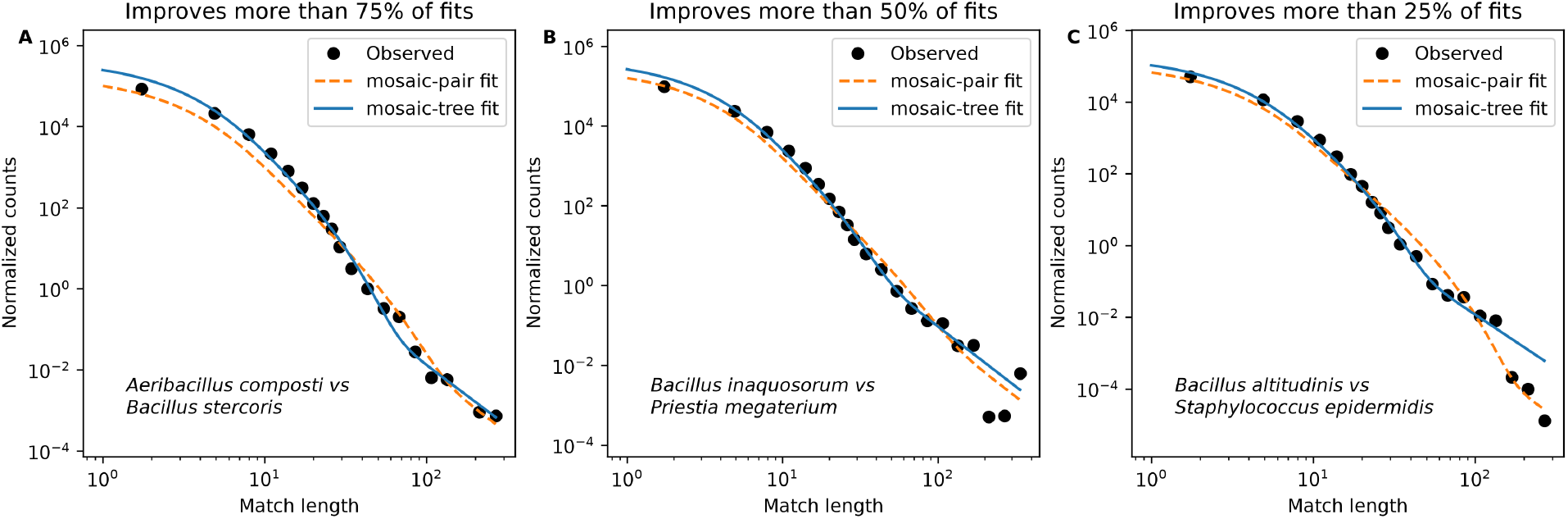
Illustration of mosaic-tree fitting improvement. Binned match length distributions (MLDs) in 3 cases (see 4.2 for plotting method). A-C: Examples of fits with `mosaic-pair` (orange dashed lines) and `mosaic-tree` (blue solid lines). They are sorted from most improved (A) to least (C), and represent the quartiles of the distribution of residual difference: 25% of fits have improved more than A, 50% more than B, and 75% more than C.

To quantitatively assess these improvements, we computed the residuals of both `mosaic-pair` and `mosaic-tree` for all pairs of species in the 3 studied clades. In *Bacillales* and *Methanobacteriota*, fits of `mosaic-pair` had very large residual values, showing that it generalized badly outside of *Enterobacterales*. In contrast, `mosaic-tree` presented similar residual values in all three clades (see also Supp. Fig. S3 and S4 for illustrations), demonstrating the broader validity of the model.

Still, we note that the residuals remain very high for some pairs (for instance *Priestia megaterium_C* and *Priestia megaterium_D*, Supp. Fig. S6). In most cases, poor fits were obtained for pairs of species that are very close, as already noted in (Sheinman et al., 2024).

### 2.4 Trees constructed with mosaic-tree are better

To study the quality of the tree inferred using `mosaic-tree`, we used the *δ* measure of treelikeness. It measures by how much a quartet of tips fails to satisfy the four-point condition (Holland et al., 2002). The closer to zero, the more treelike the quartet is. It scales to estimated distance matrices by computing *δ* in every quartet of tips.

Computing *δ* values for our three example clades, (Figure 4), we found that the distances calculated with `mosaic-tree` for *Bacillales* and *Methanobacteriota* showed a notable progress of treelikeness, from 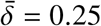 to 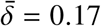 for the former, and from 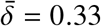 to 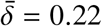 for the latter. In contrast, we found no real improvement of the inferred distances for *Enterobacterales*, from 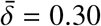 to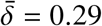, the topology also remaining identical. Note however that treelikeness is strongly influenced by the underlying tree topology (Holland et al., 2002), and is not an absolute measure, such that we do not compare the values obtained across different clades.

**Figure 4:**
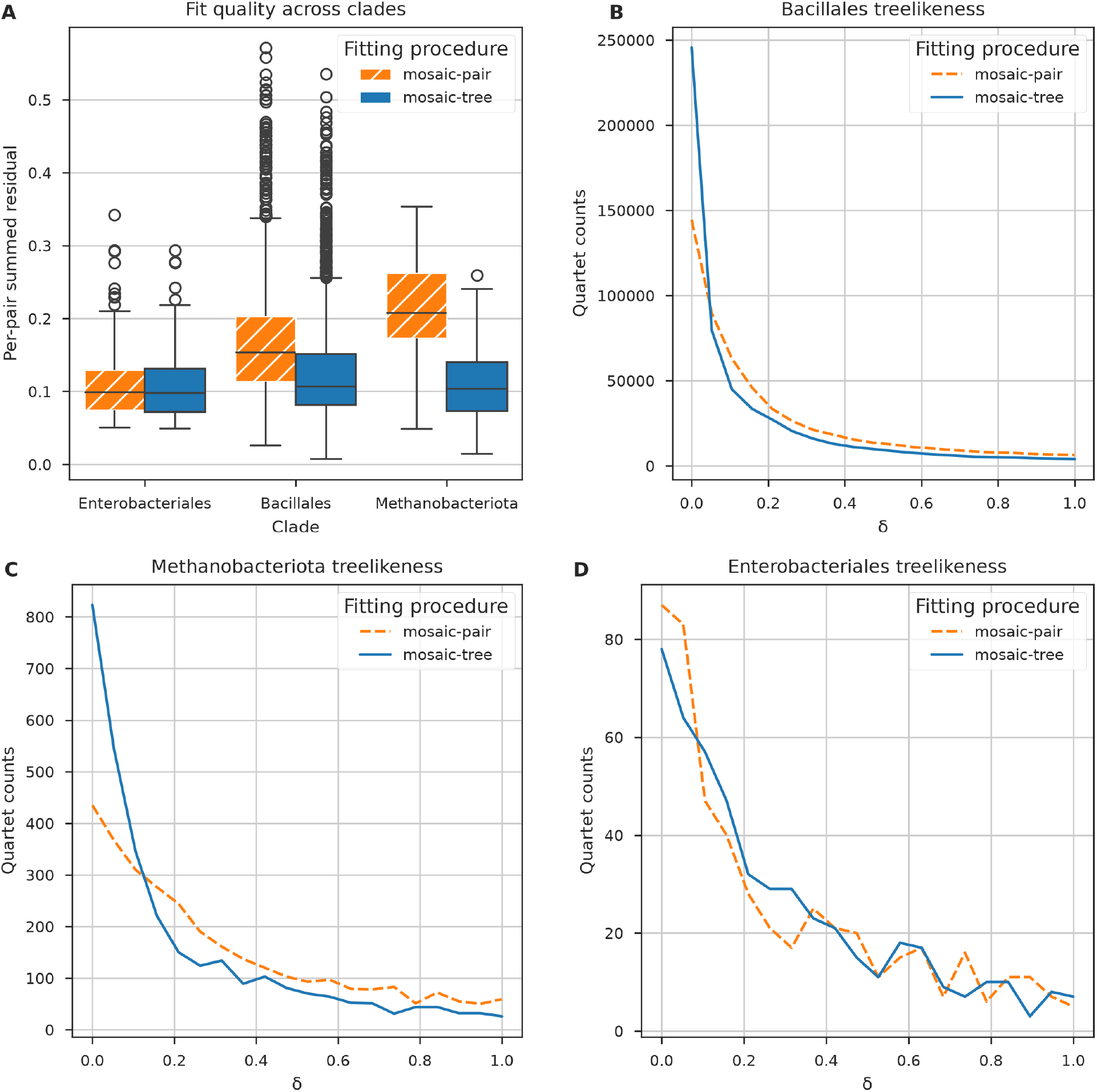
Comparison of mosaic-tree and mosaic pair model quality. **A**: Per-pair mean relative least square fit residual for all pairs of genomes belonging to all clades selected in this (*Bacillales, Methanobacteriota*) and the previous study (*Enterobacterales*). In orange hatching, the residuals for `mosaic-pair`, in solid blue, those of `mosaic-tree`. **B-D**: *δ*-plot for `mosaic-pair` (dashed orange) and `mosaic-tree` (solid blue), for *Bacillales* **(B)**, *Methanobacteriota* **(C)** and *Enterobacterales* **(D)**. The *δ*-plot is a histogram of the quartet delta value measuring treelikeness Holland et al. (2002) for every quartet of the tree.

In order to better understand whether some part of the tree were more subject to improvement than others, we also computed a per-species delta value improvement (see see Material and Methods 4.2). We found that the *δ*-values improved for all clades, showing that `mosaic-tree` was a general improvement and not specific to a given sub-tree.

Finally, we want to stress that the treelikeness is an external metric to the model’s fitting objective (minimizing residual values). The fact that both measures improve considerably and consistently can be seen as evidence, not only of the improvement, but also of the validity of `mosaic-tree`.

### 2.5 Differences in *κ* value between clades highlight differences in substitution rate distributions

In the case of *Enterobacterales*, the inferred value of *κ*, the ratio between the fastest and slowest rates, was lower than previously assumed (24.9 instead of 36.4), meaning that overall the substitution rate was less distributed than what we deduced from literature estimates Sheinman et al. (2024). We found even smaller values of *κ* in *Bacillales* (9.98), and *Methanobacteriota* (7.46).

To test whether those differences in the value of *κ* were due to a difference in the number of genomes we used in our analysis, we applied `mosaic-tree` to *Enterobacterales* and *Bacillales* with a lower number of genomes (Supp. Fig. S7, S8 and Supp. Table. S1). The estimated values of *κ* remained stable, showing that the differences we observe between clades do not depend on the number of genomes used in the inference.

### 2.6 Trees can be calibrated or scaled

When using the previously defined 3.64 × 10^−9^bp^−1^yr^−1^ as the upper bound of the substitution rate distribution, we can scale the tree to time. Indeed, this quantity can be relatively easily estimated via mutation accumulation experiments (Lenski et al., 1991). However, as 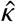 was lower than assumed in our previous study, the time value estimated for species divergences changed. For instance, we obtained a time divergence between *Escherichia coli* and *Salmonella enterica* of only about 50 Myrs, much smaller than the commonly accepted 140 Myrs (Ochman and Wilson, 1987), and half of `mosaic-pair`’s previous result. This discrepancy could be explained if the top substitution rate was lower than assumed here.

Another approach is to fix the upper bound of the rate distribution through commonly accepted time divergences. For instance in the case of *Enterobacterales*, using the aforementioned divergence as a fixed point, the highest substitution rate was 1.32 × 10^−9^bp^−1^yr^−1^ and the lower 5.30 × 10^−11^bp^−1^yr^−1^. This top substitution rate of the distribution was not far from three times smaller than measured in (Wielgoss et al., 2011), the reference used in our previous work. One explanation could be that over long periods (the smallest time divergence of the *Enterobacterales* tree, using 3.64 × 10^−9^bp^−1^yr^−1^ as the maximum substitution rate, was 12.8 Myrs between *Escherichia coli* and *Escherichia fergusonii*), all segments of either genome have been under selection at some point, and the highest substitution rate was lower than a background mutation rate measured in much shorter evolution experiments.

### 2.7 Inferred Phylogenies

We first examined whether `mosaic-tree` was able to recover the results obtained by `mosaic-pair` on *Enterobacterales* in our previous study. Apart from the previously mentioned time discrepancies when using the same calibration method, we found the same topology with `mosaic-tree` and with `mosaic-pair`. There are two differences between the GTDB topology and the two `mosaic-tree`/`mosaic-pair` topologies (Supp. Fig S10), discussed in Supp. Mat. 10.1.

#### 2.7.1 Bacillales

*Bacillales* are an interesting case study as their phylogeny has been updated recently (Gupta et al., 2020). As with `mosaic-pair`, once the pairwise time divergence estimate were collected, an UPGMA (Sokal et al., 1958) tree was computed (Figure 5).

**Figure 5:**
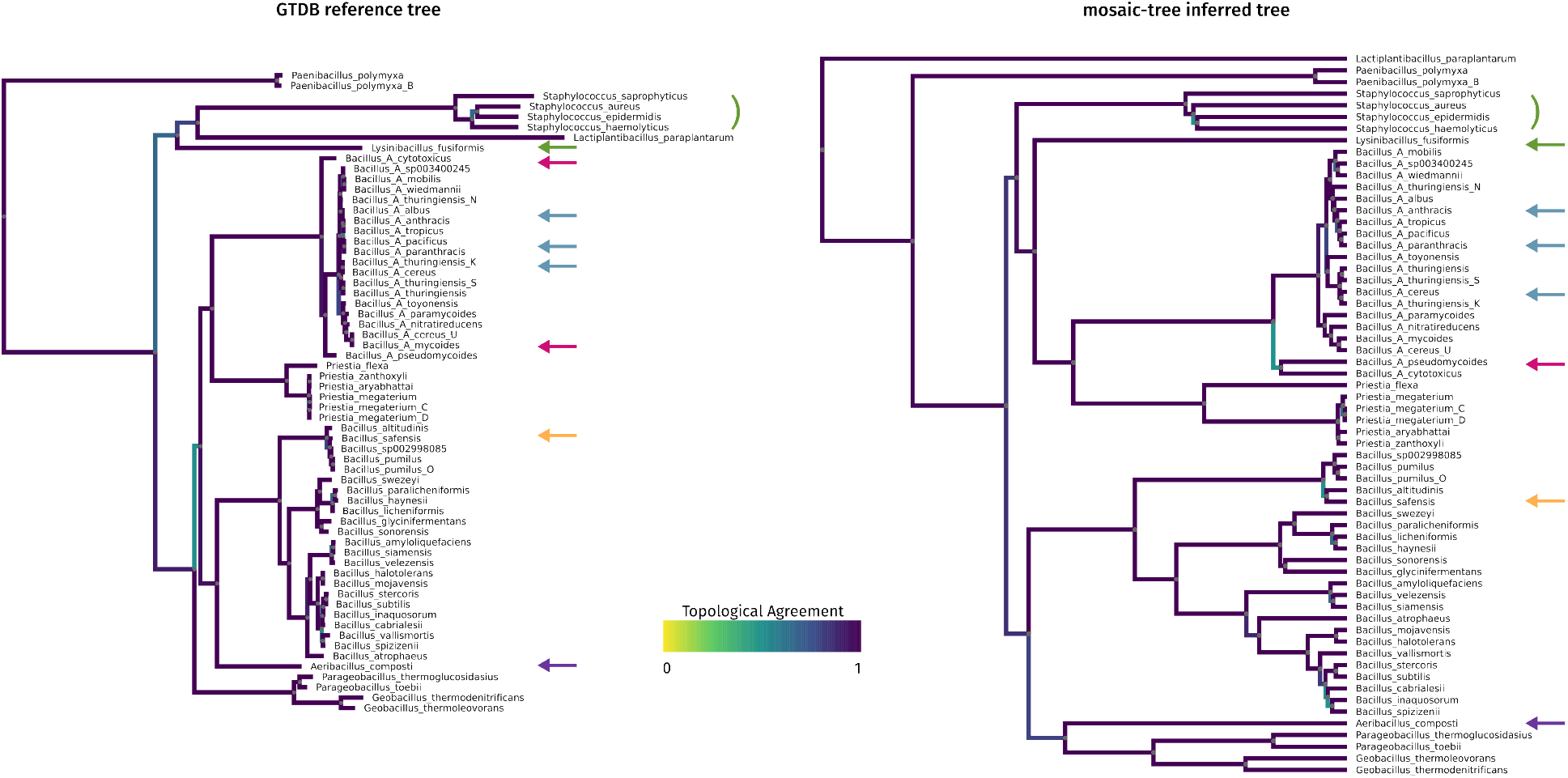
Reference *Bacillales* tree and estimated *Bacillales* tree. Reference tree from the GTDB database (left) and inferred tree by the `mosaic-tree` method, for the *Bacillales* and selected outgroups (*Paeanibacillales, Lactiplantibacillus paraplantarum, Staphylococcales, Lysinibacillus*). Branches are colored by topological agreement. Arrows indicate specific species referenced in the text: the placement of outgroups *Lysinibacillus fusiformis* and *Staphylococcales* (green), the grouping of *Bacillus_A cytotoxicus* and *Bacillus_A pseudomycoides* (pink), the correct separation of *Bacillus_A paranthracis* and *Bacillus_A cereus* (blue), the grouping of *Bacillus altitudinis* and *Bacillus safensis* (yellow) and the placement of *Aeribacillus composti* (purple).

There was agreement between the GTDB tree (used here as a reference) and our inferred tree (Figure 5), both in relative lengths of branches and clustering information. Some of the selected outgroups (species of *Paenibacillales, Lactobacillales*) were recovered as outgroups in our inferred tree. On the other hand, *Bacillales_A*’s *Lysinibacillus fusiformis* (green arrow on Figure 5) and selected species of *Staphylococcales* (green brace) were not outgroups in `mosaic-tree`’ species tree, but rather sister clades of *Bacillaceae_G*. The placement of *Lysinibacillus fusiformis* is an interesting disagreement, as it has changed order in different releases of GTDB, being a member of *Bacillales* in releases 95 and 214 while being in its own order, *Bacillales_A*, in current releases. The placement of *Staphylococcales* is somewhat more puzzling even though only a short branch separates the nodes leading to this group and separating *Bacillales* in *Bacillaceae_G* and *Bacillaceae*. Indeed, in GTDB, *Staphylococcales* have always been a separate group from *Bacillales*. In the NCBI taxonomy however, *Staphylococcacae* are part of *Bacillales*. We investigated in more details by iteratively removing all possible combinations of the four *Staphylococcales* species (Supp. Fig. S12 shows two examples). It revealed that *Staphylococcus saprophyticus* and *Staphylococcus epidermidis*, alone or in combination were placed outside the *Bacillaes*, whereas this was not the case for *Staphylococcus aureus* and *Staphylococcus haemolyticus*. This is also noticeable on a heatmap of the inferred distances (Supp. Fig. S13), where *Staphylococcus aureus* and *Staphylococcus haemolyticus* are closer to some species of *Bacillaceae_G* than to the other two *Staphylococcales*. Overall the placement of *Staphylococcales* as a group is quite unstable with our model.

For the family *Bacillaceae* we note that the first disagreement, the placement of *Bacillus altidudinis* and *Bacillus safensis* (orange arrows on Figure 5), appeared three nodes away from their most recent common ancestor (MRCA). It indicates that disagreements remain contained to the relative placement of distal nodes. For *Bacillaceae_G*, the second important family of *Bacillales*, disagreement was notable higher in the tree, as *Bacillus_A cytotoxicus* and *Bacillus_A pseudomycoides* (pink arrows on Figure 5) form a cherry in the tree inferred by `mosaic-tree`, but not in the reference. The branch lengths between the consecutive inner nodes leading to *Bacillus_A cytotoxicus* and *Bacillus_A pseudomycoides* were short. Finally, *Aeribacillus composti* (purple arrows on Figure 5) was grouped with the *Geobacillus* species, and together this group was placed as a sister group of family *Bacillaceae*. While most differences involved short inner branch lengths on both trees, therefore carrying low significance, the placement of *Geobacillus* and *Aeribacillus composti* was a more substantial difference.

In particular, in agreement with recent classification changes (Liu et al., 2017), *Bacillus_A anthracis* and *Bacillus_A paranthracis* clustered together, contrary to previous classification that considered the latter as part of *Bacillus_A cereus*. `mosaic-tree` was able to correctly place the two species using the entire genome sequences, without gene annotation and filtering or multilocus sequence typing (as was done in (Liu et al., 2017)). This was achieved even though parts of the genomes are discordant with this classification, as evidenced by the fact that using 16S sequences is not enough to classify genomes of this group into species (Liu et al., 2017).

#### 2.7.2 Methanobacteriota

For *Methanobacteriota*, MLDs were comparable to the ones from *Bacillales*, and to *Enterobacterales* (see Supp. Fig. S5). The range of the power law behavior was shorter than on *Bacillales*, which was consistent with the fact that the divergences were less distributed (*κ* is smaller).

We obtained the tree visible in Figure 6. Overall disagreement with GTDB reference tree was more important than in the case of *Bacillales*, which can be expected as less genomes are available in databases, and as *Archeae* phylogeny is less established. Comparing with the reference tree from GTDB, the monophyly of *Methanobacteriaceae* was not recovered. Indeed, *Methanothermobacteriaceae* (green brace on Figure 5), a different family, was in our case placed inside of *Methanobacteriaceae*. Support for the placement of the ancestor node of *Methanobacteriaceae* and *Methanothermobacteriaceae* is however low in the reference tree, which suggests that the current reference phylogeny is not definitive. At the genus level, agreement was higher: indeed, *Methanobacterium, Methanothermobacter* are identical, and *Methanobrevibacter_A* and the other *Methanobrevibacter* were similar between our inferred tree and GTDB’s reference. A notable disagreement was the placement of human commensal *Methanosphaera stadtmanae* (green arrow on Figure 5) as a sister clade of *Methanobrevibacter* in our inferred tree, while it is within the *Methanobacterium* genus in the reference.

**Figure 6:**
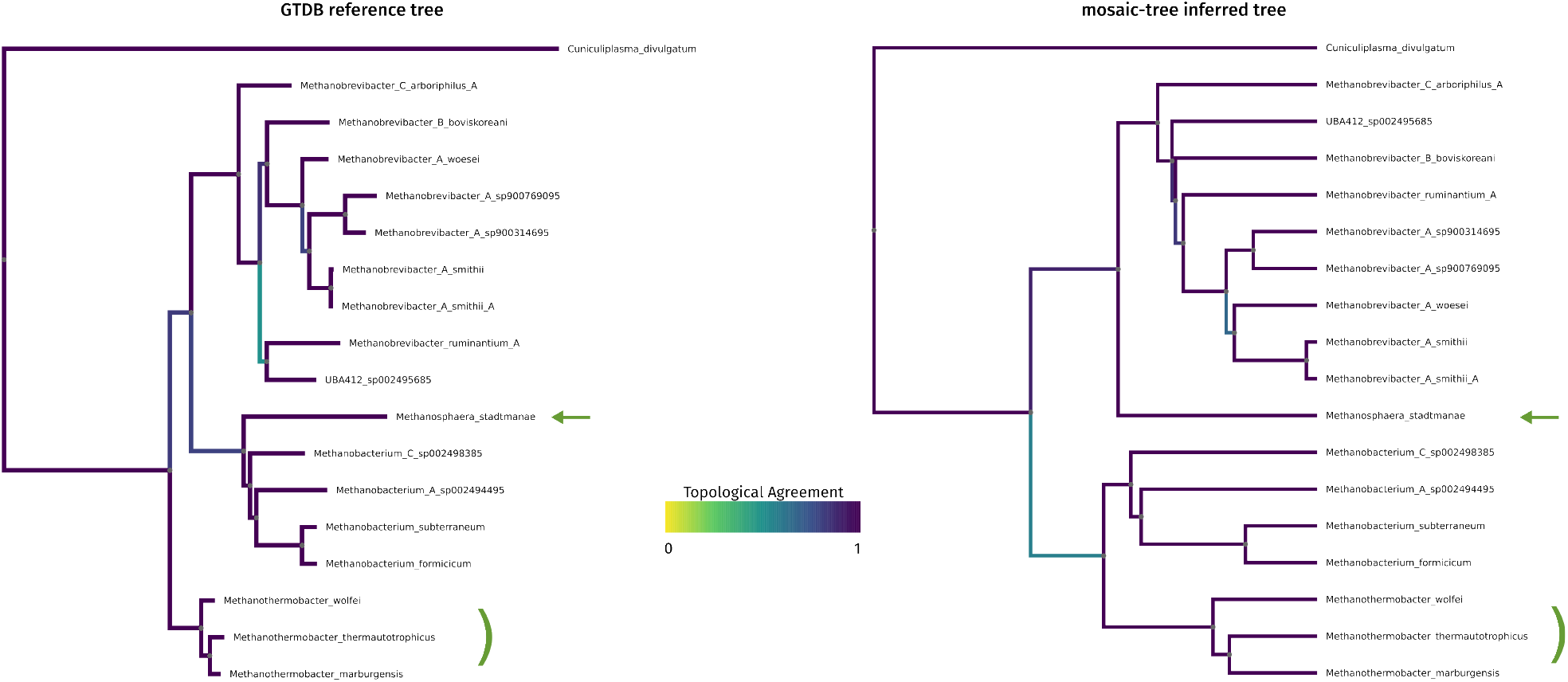
Reference *Methanobacteriota* tree and estimated *Methanobacteriota* tree. Reference tree from the GTDB database (left) and inferred tree by the `mosaic-tree` method, for the *Methanobacteriota*, with *Cuniculiplasma divulgatum* as an outgroup. Branches are colored by topological agreement. The green arrow highlights the disagreement in placement of *Methanosphaera stadtmanae* and the green brace the disagreement in the placement of *Methanothermobacteriaceae*.

## 3 Discussion

In this study, we propose `mosaic-tree`, a method to estimate a time divergence and HGT rate from whole genome alignments. `mosaic-tree` builds on principles established by our previous study (Sheinman et al., 2024), and allows inference in deeper evolutionary times and across a broader range of taxa, exemplified here in *Bacillales* and in the Archeae phylum *Methanobacteriota*. To achieve this, we used a more sensitive aligner, and fitted the bounds of the substitution rate distribution, relaxing the previous model. Together, those changes allowed our `mosaic-tree` method to infer trees for complex clades, with good topological agreement with the literature. Overall, our results demonstrate that this method is applicable to a wide range of species, hinting at the universality of the underlying model.

The main novelty of `mosaic-tree` is the addition of a free parameter to the model, *κ*, which controls the width of the substitution rate distribution. This single additional free parameter both substantially improved the fit of the model and the treelikeness of the inferred distance matrices. Notably, this parameter was different for each of the studied clades. Whether the variation in *κ* has biological or technical origins remains to be investigated: indeed, species sampling, tree depth or topology could influence the value of *κ*. Therefore the elucidation of the link of this parameter to evolutionary processes is left to further studies. More broadly, the assumption of one distribution of rates for every comparison might be oversimplifying. Dropping the assumed ultrametricity of the inferred distances, it could be possible to fit one *κ* parameter per comparison, taking the median of the distribution as a distance, and using Neighbor-Joining (Saitou and Nei, 1987) to compute the tree. This could improve the suboptimal model fits for close species as well as the tendency of the fit to compensate a decreased *κ* by an increase in the transfer rate.

The `mosaic-pair` and `mosaic-tree` model assume that the distribution of the substitution rates is linear. This is not straightforwardly explainable through the sum of two stable rate distributions of a pair of aligned genomes (see Supp. Mat. 10.3), but could arise through completely correlated and linear rate distributions in all the considered species. Alternatively, rate evolution could induce approximately linear rate distributions. The first case is biologically very restrictive, but could be a reasonable approximation for very conserved essential genes. On the other hand, rate evolution is more flexible, but it is not self-evident that there are processes whose steady state is an approximately linear distribution of rates. In this study we simulated trees, as opposed to only pairs of genomes in our previous study, to investigate `mosaic-tree` in the two aforementioned scenarios. We found that tree topology was recovered in both cases, even though distances are underestimated when we simulate rate evolution with a bounded Brownian motion. Strict adhesion to the linear rate distribution is thus unnecessary to infer tree topologies, and the linear approximation is a useful approach.

Despite those limitations, the tree topologies obtained with `mosaic-tree` are similar to those of the literature, showing the applicability of the method. Comparing inferred trees to the reference ones, we found that short branches are shorter in the reference than in `mosaic-tree` trees. One explanation for this difference could be that aligning whole genomes results in a flexible alignment length, dynamically leveraging the amount of shared information between each genome pair, in contrast to classical methods that select solely widely shared genes across a taxon.

Time-calibrating our trees is possible, either by fixing the upper bound of the rate distribution, or by fixing a specific branch and scaling the rest of the tree with it. However it leads to new questions. Using the same upper bound as in our previous study, sourced from (Wielgoss et al., 2011), the estimated time divergences for *Enterobacterales* became almost three times smaller than the estimates from the literature. It might indicate that using the background substitution rate to calibrate time estimates is overestimating the number of substitutions of the loci under the lowest selective pressure. To a lesser extent, the same trend was apparent in simulations with a rate evolution, outside of the model hypotheses. Therefore a promising way to investigate this underestimation would be to incorporate rate evolution into our modeling.

To this end, we would need to refine the linear rate distribution approximation, and potentially relax the distribution shape assumption, but it would require more statistical power to fit more parameters. In this study, we considered the set of pairwise distances to infer a tree-wide parameter. But it would be feasible to use multiple genome alignment and their distribution of match length, as exemplified by our study of HGT networks in *Enterobacterales* (Sheinman et al., 2026), to further improve `mosaic-tree`. Using a rate evolution model, we could leverage the information of multiple unique matches (efficiently available with tools such as mumemto (Shivakumar and Langmead, 2025)) to better estimate evolutionary parameters. This could potentially provide additional constraints for the inference.

Overall, we demonstrate that `mosaic-tree` has a wide applicability, allowing for distance estimation using whole genomes of Bacteria and Archeae with few hypotheses. Our results highlight the importance of substitution rate distributions, and that information on those is contained in MLDs. With an adequate model of substitution rate evolution, our model would have the power to improve bacterial tree calibration, a promising avenue for further study.

## 4 Materials and Methods

### 4.1 Data selection

We used the Genome Taxonomy Database (Parks et al., 2022) release r226 both as a reference and for taxonomic definitions. In this study, all taxonomic names refer to the GTDB classification.

For *Bacillales*, we selected all species with at least 10 genomes in GTDB, a completeness of above 90% and contamination under 5% according to checkM2, the N50 of contigs being above 100kb and that of scaffolds above 150kb. We also enforced the presence of at least 18 tRNA genes coding for different amino acids. In the end we capped the number of genomes per species at 50, ending up with 2357 genomes. This amounted to on average 38 genomes per taxa and 2,725,961 alignments.

For *Methanobacteriota*, we selected all species with at least 5 genomes and *Thermoplasmataceae* as an outgroup, ending up with 200 genomes. This amounted to on average 11 genomes per taxa and 17,485 alignments.

For *Enterobacterales*, we selected clades in the same manner as our previous study (Sheinman et al., 2024) but decreased the amount of genomes used, for fairer comparisons and to save on compute time. We then filtered the genomes completeness of above 90% and contamination under 5% according to checkM2, and less than 50 contigs. We also enforced the presence of at least 18 tRNA genes coding for different amino acids. In the end we capped the number of genomes per clade at 200, ending up with 1743 genomes. This amounted to on average 145 genomes per taxa and 1,366,301 alignments. Accessions for the assemblies used here for analysis are available in supplementary materials.

### 4.2 Inference Pipeline

We used lastz (Harris, 2007) for whole genome alignment. To eliminate mobile genetic elements that are not taken into account in the model, all genomes were self-aligned and all repeats are N-masked. Then genomes were aligned with lastz using default parameters.

To represent and fit MLDs, we proceeded similarly to our previous study (Sheinman et al., 2024), namely we used a linear 3 bp binning up to 30 bp and logarithmic binning for larger matches: [1, 3], [4, 6], …, [28, 30], [31, 30× 10^0.1^], [30×10^0.1^ +1, 30×10^0.2^]… In the plots, centers of the bins are geometric mean of their boundaries.

Within each bin, we count the number of matches and normalize it by the size of the bin. In addition, we normalize the MLD by the total number of alignments we do for the two considered taxa. If we analyze two taxa with n1 and n2 genomes, respectively, we do n1 × n2 alignments, collect all the exact matches, and, therefore, divide the total MLD by n1 × n2.

From empirical MLDs, acquired through whole genome alignment of multiple pairs of genomes from each species, a transfer rate and a distance were co-estimated. Estimation is made by fitting binned empirical MLDs to the model using mean square relative difference. All pairs of species were fitted together, each comparison with two parameters, the distance and the relative HGT rate. In addition we fitted *κ*, the ratio between the fastest and slowest rates of the distribution. Optimization started by grid scanning for the best starting value of *κ*, keeping it fixed and fitting each species pair *i*’s parameters, *θ*_*i*_ and *ξ*_*i*_ (respectively the evolutionary distance and the relative horizontal transfer rate, see also 10.4) using the L-BFGS-B algorithm (Byrd et al., 1995; Zhu et al., 1997). The three best starting point were refined through block coordinated descent, iterating on fitting *κ* alone and the pairwise parameters *θ*_*i*_ and *ξ*_*i*_ alone until convergence. Each of them was finalized by a last run of L-BFGS-B, optimizing all model parameters (pairwise *θ*_*i*_, *ξ*_*i*_ and *κ*). The best result was retained.

Treelikeness is computed with the `delta.plot` function of the ape package (Paradis and Schliep, 2019). In particular, the species-wise progress in treelikeness is the relative difference in 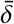 (averaged over all quartet of species a given species participates in) between `mosaic-pair` and `mosaic-tree` distances.

The estimated distances describe an ultrametric tree, as the underlying summed substitution rate distribution is assumed to be shared by all pairs of bacterial species. UPGMA (Sokal et al., 1958) was thus used for computing the tree. Tree representations for inferred trees and reference trees were drawn with Phylo IO (Robinson et al., 2016). The degree of similarity, indicated in colors for branches, is based on the Jaccard index (Robinson et al., 2016).

The rest of the pipeline is written in python, allows for parallelization, and uses a SQLite database for the storage of the match length distributions. The code base is available on https://github.com/CBIO-mines/mosaic_me

### 4.3 Simulation pipeline

A species tree was either fixed or simulated with the implementation of the Birth Death model in ape (Paradis and Schliep, 2019).

We simulated genomes with piecewise constant mutation rates according to the two models mentioned in 2.2, the first one with no rate evolution, each orthologous segment having the same substitution rate, and the second where rates evolve following a bounded random walk. For the first case, gene trees were scaled by a rate drawn from a linear distribution. For the bounded random walk, the root substitution rate was drawn from a uniform distribution *U*(*µ*_*min*_, *µ*_*max*_), and then the rate walked between *µ*_*min*_ and *µ*_*max*_ for a given number of steps (5 × 10^4^ for section 2.2) for the length of the tree (a distance of 4 × 10^8^ scaled with *µ*_*max*_ = 1). For every branch, the rate was computed as the average of the ascendant and descendant nodes’ rates as has been done previously (Kishino et al., 2001; Lartillot et al., 2016). The process was repeated for *N* gene trees, and each gene tree was scaled by the root age. Gene tree manipulation was done using skbio.tree (Aton et al., 2026). We used Alisim (Ly-Trong et al., 2023) with a fixed sequence length to generate sequences. Resulting gene sequences were concatenated to emulate the gene-rich genomes of bacteria. Distances indicated in results were scaled with *µ*_*max*_ = 1.

In order to simulate MLDs directly from gene trees, for Supp. Fig. S2, the number of substitution of a gene alignment was drawn from a Poisson distribution with parameter *λ* = substitution rate × branch length × gene length. The position of the substitutions were then uniformly drawn in the gene. The code used to generate the simulation of this paper is available at: https://github.com/CBIO-mines/sim_alisim.

### 4.4 Aligner sensitivity

The increased resolution of the `mosaic-pair/tree` compared to other distance metrics comes in part from modeling the sensitivity of the aligner: we assume that above a certain sequence divergence, it cannot align sequences anymore. Thus the collection of matches that we observe comes from a truncated version of the distribution of divergences. In order to improve on `mosaic-pair`, we replaced MUMmer (Marçais et al., 2018) with lastz (Harris, 2007), trading off compute time for sensitivity. Therefore, we needed to update the cutoff of the distribution. Previously, we assumed it to be truncated above *δ*_*al*_, *δ*_*al*_ being the maximum distance recovered by the aligner — the proportion of mismatches. However this severely underestimated *θ*: indeed, a proportion of the bases aligned would have mutated multiple times in either of the sequences. Using the classical evolutionary model JC69 (Jukes and Cantor, 1969), we corrected for unseen substitutions, changing the value of *δ*_*al*_ from 0.5 to 0.82. We discuss and measure the impact of this change in Supp. Mat. 10.2. This correction on the aligner’s sensitivity was used in all estimations in the present study.

## 7 Data availability

The data underlying this article, genome assembly accessions, are available in Supplementary Materials.

## 8 Author contributions statement

P.E. wrote the software, curated the data, conducted the experiments, analyzed the results and designed the visualizations, A.H. curated the Archeae data, conducted and analyzed the Archeae results, M.S., F.M, P.A., T.S. and P.E. conceived the experiments and the methodology, F.M. and C.A. supervized P.E., P.A. and F.M. acquired resources and compute, P.E. wrote the draft manuscript, M.S., F.M, P.A., T.S., C.A.. A.H. and P.E. edited and reviewed the manuscript.

## 9 Acknowledgments

Claude Code (Opus 4.6, Sonnet 5) was used for three separate and clearly delineated tasks: i) code review and documentation once all the code was written, ii) to develop the optimization procedure to fit the model simultaneously across all pairwise comparisons, iii) light proofreading of the manuscript.

