## Supplementary for "Inferring evolutionary distances from whole prokaryotic genome alignments, accounting for among-locus substitution rate heterogeneity"

### 10 Supplementary

#### 10.1 Differences between GTDB and `mosaic-tree`/`mosaic-pair` *Enterobacterales* trees

There are three differences between the GTDB and `mosaic-tree` and (Sheinman et al., 2024) trees. The position of *Escherichia albertii* and *Escherichia fergusionii*, as sister species to *Escherichia coli*; in addition *Salmonella enterica* and *Citrobacter* form a cherry in the `mosaic-tree` countrary to the GTDB phylogeny. Finally, *Klebsiella pneumoniae* goes from being a sister clade to the *Escherichia*, *Salmonella enterica* and *Citrobacter* group to being a sister clade of *Enterobacter*. Once again however, all involved branch length are small on the GTDB tree.

#### 10.2 Jukes-Cantor correction tests

In order to test whether the `mosaic-pair` model was applicable outside the *Enterobacterales* clade, we tested it on *Bacillales*. Applying `mosaic-pair` failed because of the low sensitivity of the aligner (MUMmer) used in the previous version (Supp. Fig. S14). Indeed, for the same set of genome comparison, lastz was capable of retrieving many more alignments (See Supp Fig. S11). In the case of a very sensitive aligner such as lastz (Harris, 2007), the fit was strongly improved if we corrected the aligner sensitivity with an evolutionary model, as was verified in simulations (Supp. Figure S15).

#### 10.3 Underlying substitution rate distributions

In `mosaic-pair`, the rate evolution in each species that were compared was hypothesized to be uniformly distributed, and i.i.d. This was the a potential justification for the linear distribution of their sums, and the $-4$ power laws observed in the data. However, this assumption is incompatible with a tree representation of genome evolution. Indeed, in the case of a species tree of 3 species A, B and C, each pair cannot consist of only a sum of two substitution rates (Fig. S1). Either comparisons to the outgroup involve a third substitution rate between the root and the cherry, or the cherry and the branch to the root have only one substitution rate. In either case assumptions made by the model are not met, while comparing AC and BC, or AB. The first case involves summing 3 i.i.d. substitution rate distributions, which, for reasons developed in (Massip et al., 2016) would imply a power law of exponent $-5$ for the vertically transmitted part. The consequence of complete correlation is dependent on the shape of the underlying rate distribution in the genome, but for the simple uniform distribution of rates the consequence would be a $-3$ power law, and for a linear distribution a $-4$ power law, which was used for simulations for Figure 2 (blue) and Supp. Fig. S2.

#### 10.4 Dimensionless units

In this work, we re-scaled the problem in dimensionless units: for pair $i$, we studied $\theta_i$, the expected number of substitutions per site accumulated at the background (fastest) rate $\mu_s$, so that $\theta_i = \mu_s \tau_i$ (called distance for simplicity), the ratio $\xi_i$ between HGT rate and background substitution rate (called relative HGT rate) and the ratio between the upper and lower bounds of the substitution rate distribution $\kappa$. This way, we separated inference (fitting the parameters) and calibration (choosing the upper bound of the substitution rate distribution, or fixing a pairwise $\tau$), which could be done afterwards.

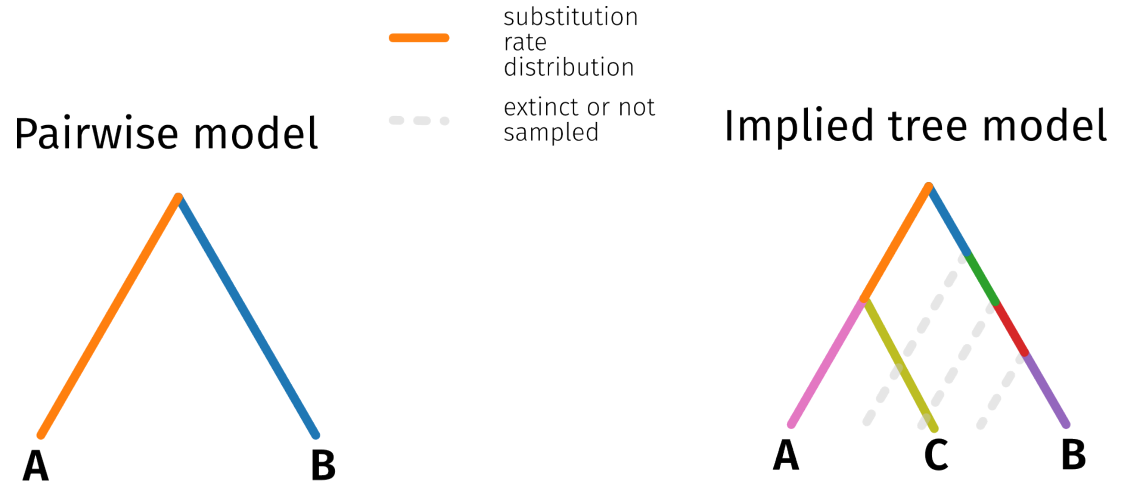


**Supplementary Figure S1: Incompatibility of `mosaic-pair` with a tree.** The summed substitution rate distribution was assumed to arise from a sum of two i.i.d. distributions for each pair in (Sheinman et al., 2024). But considering trees implies considering sums of more than two distributions.

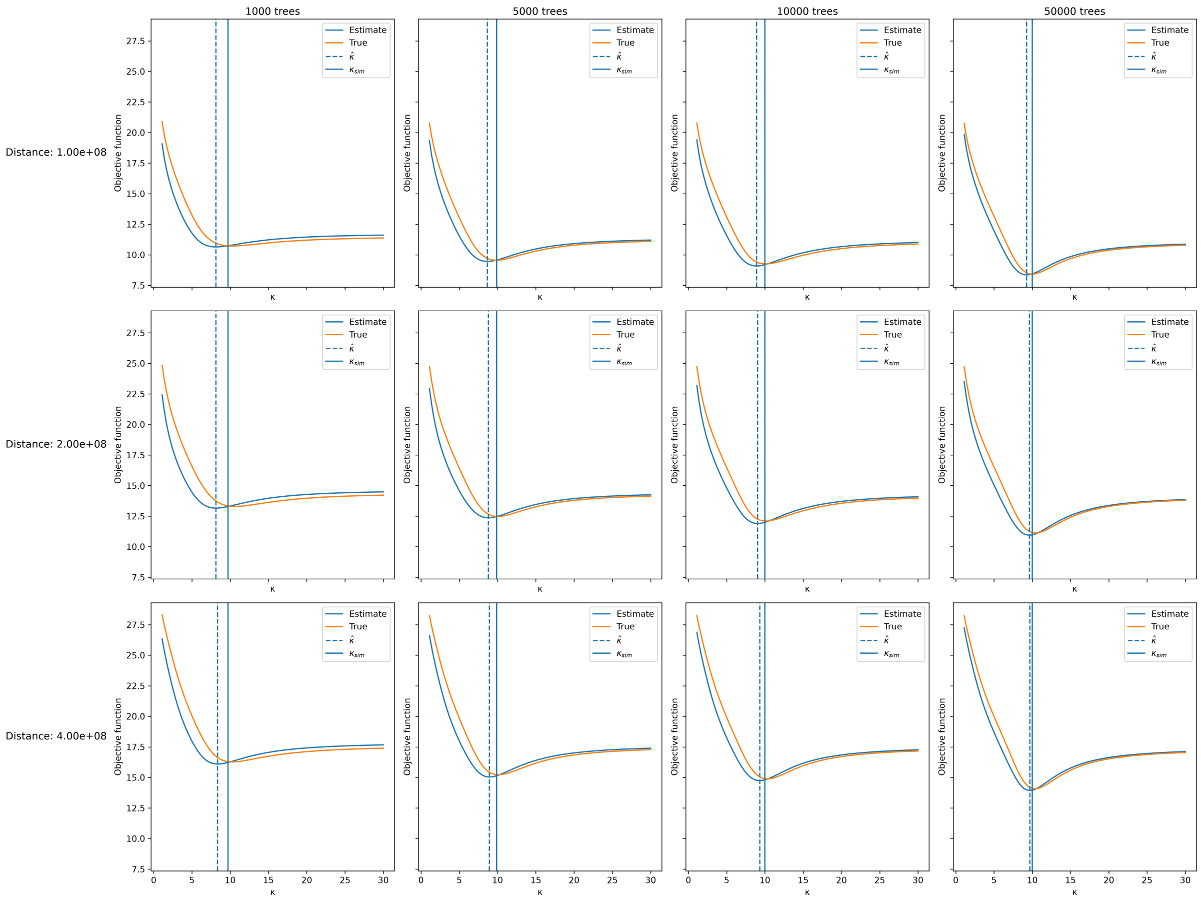


**Supplementary Figure S2: Testing the inference of simulated $\kappa$ in different scenarios.** Summed mean relative least square objective function value ($y$) at different values of $\kappa$ ($x$), for multiple tree heights (rows) and gene tree amounts (columns), with $\kappa_{\text{param}} = 10$. $\kappa_{\text{sim}}$ can be lower than $\kappa_{\text{param}} = 10$, as the minimum possible mutation rate might not be realized, depending on the number of gene trees. The same random tree is used for all simulation cases. The orange curve represents the objective function evaluated at the true parameter values (all parameters fixed at their simulated values, $\kappa$ varying). The blue curve represents the objective function evaluated at the estimated parameter values (all parameters fixed at their L-BFGS-B estimates, $\kappa$ varying). The solid vertical line indicates the true $\kappa_{\text{sim}}$; the dashed vertical line indicates the estimated $\hat{\kappa}$.

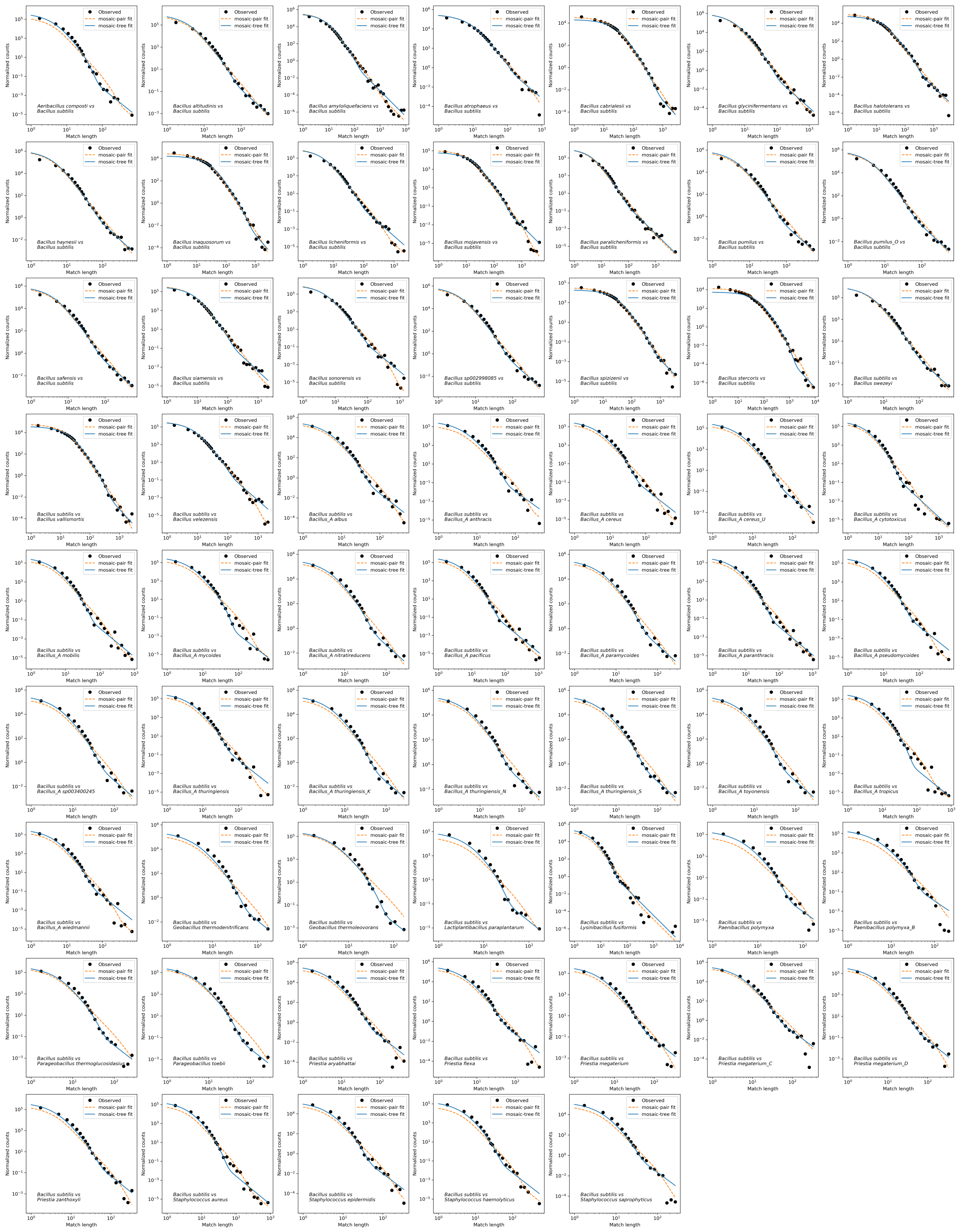


**Supplementary Figure S3:** ***Bacillales* MLD examples.** All pairwise MLDs involving *Bacillus subtilis* in the *Bacillales* clade, with `mosaic-pair` fit (dashed orange) and `mosaic-tree` fit (solid blue)

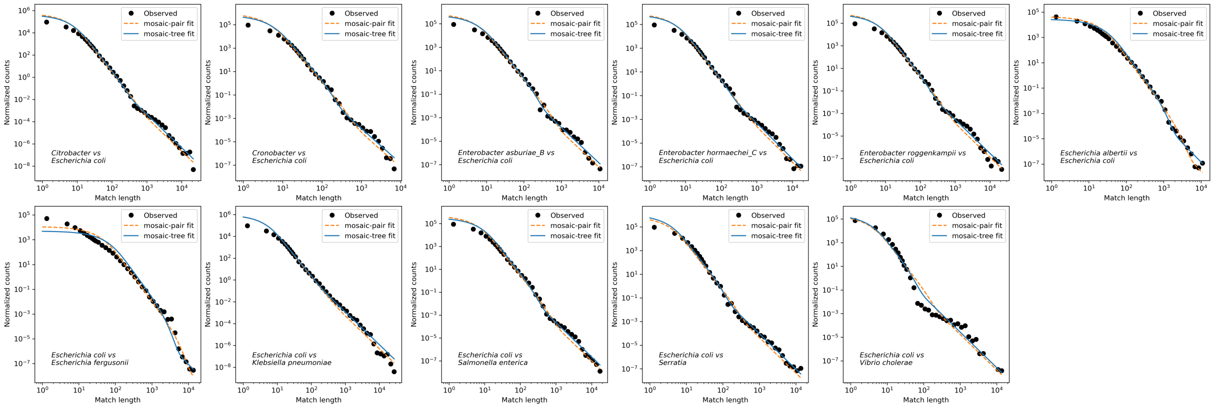


**Supplementary Figure S4:** ***Enterobacterales* MLD examples.** All pairwise MLDs involving *Escherichia coli* in the *Enterobacterales* clade, with `mosaic-pair` fit (dashed orange) and `mosaic-tree` fit (solid blue).

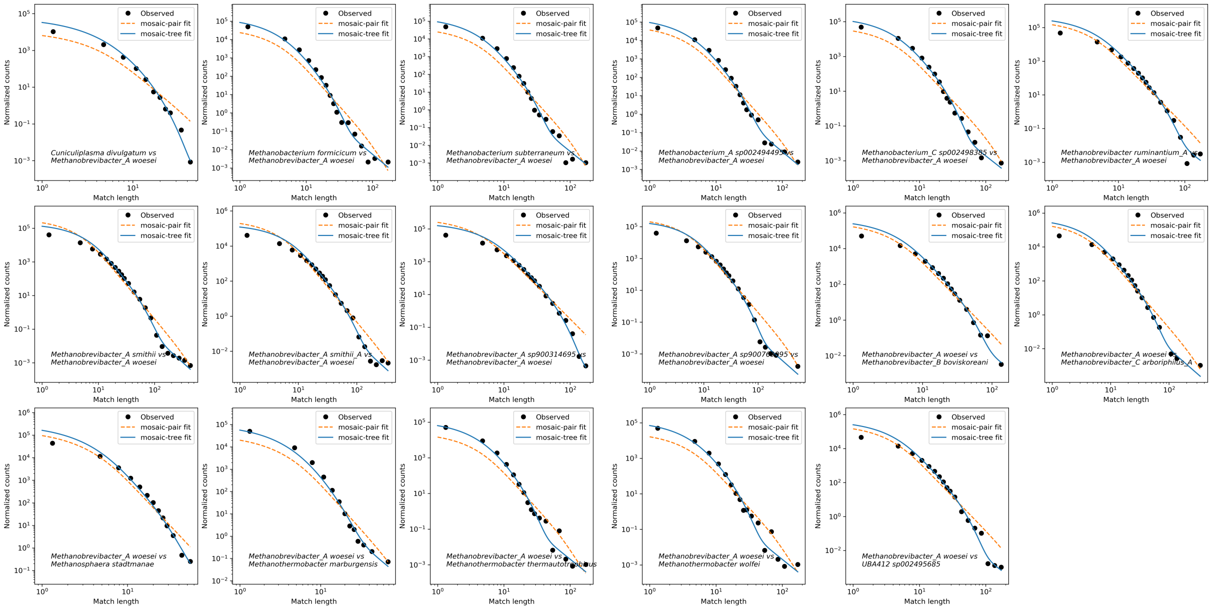


**Supplementary Figure S5:** ***Methanobacteriota* MLD examples.** All pairwise MLDs involving *Methanobrevibacter_A woesei* in the *Methanobacteriota* clade, with `mosaic-pair` fit (dashed orange) and `mosaic-tree` fit (solid blue).

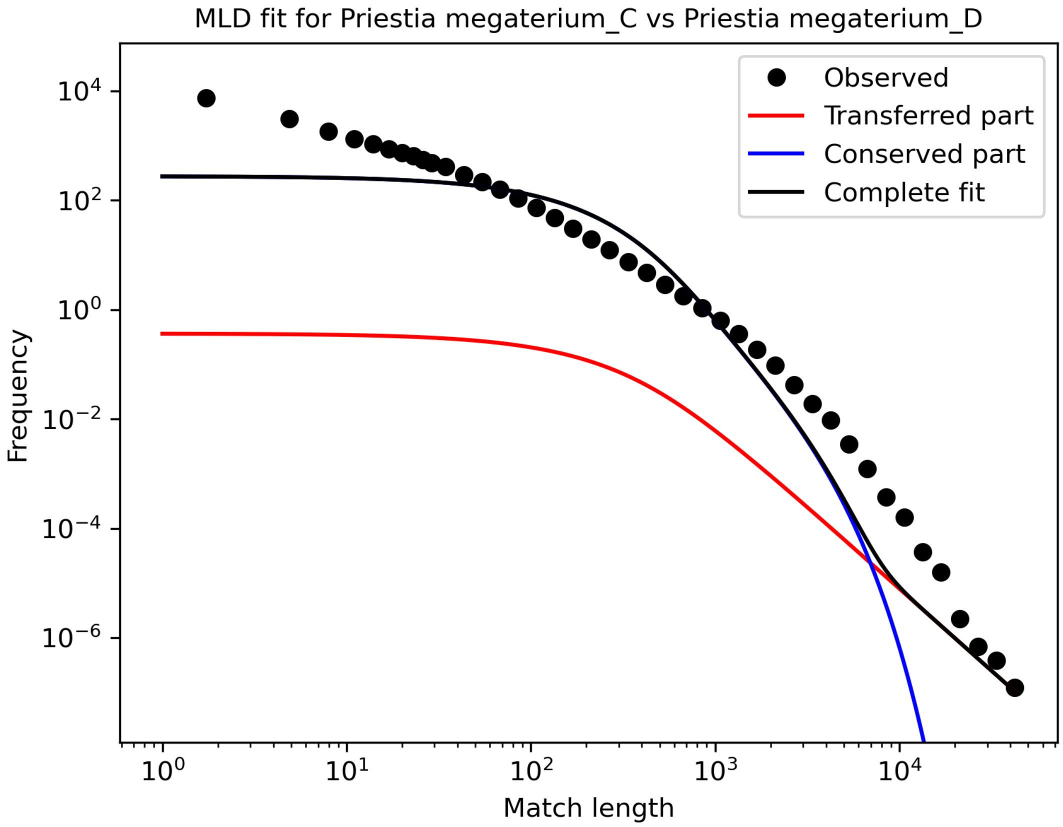


**Supplementary Figure S6:** ***Priestia megaterium_C*** **and** ***Priestia megaterium_D*** **MLD and** **`mosaic-tree` fit.** The two components of the model, the conserved, vertical component ($m_c$), a $-4$ power-law, in blue, and the HGT $m_h$ component, a $-3$ power-law, in red, are separated.

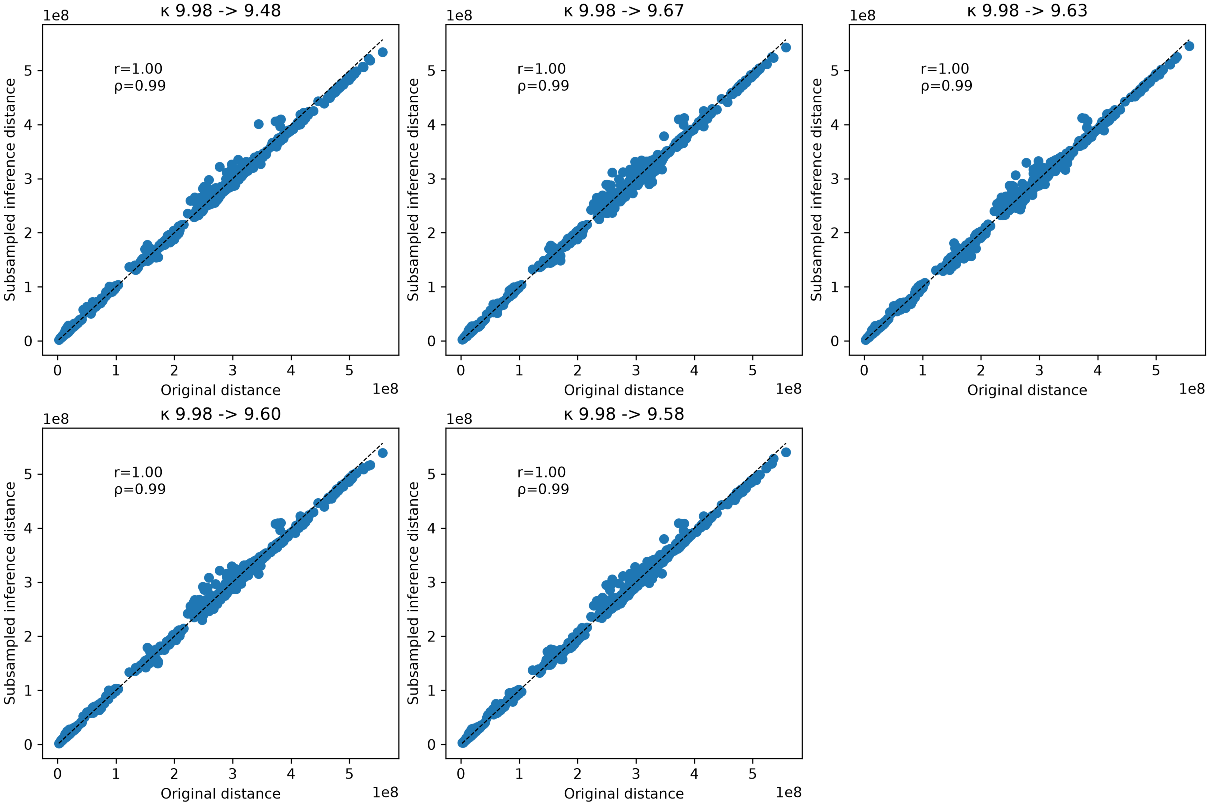


**Supplementary Figure S7: $\hat{\kappa}$ robustness to the amount of data.** *Bacillales* genomes are downsampled to the amount of data of Methanobacteriota. Each scatterplot shows the original distance inferred with full data and the distance inferred with subsampled data, along with Pearson and Spearman correlations. $\hat{\kappa}$ before and after subsampling are indicated in scatterplot titles. Each plot is one of five repeats to judge robustness.

| Clade | $\hat{\kappa}$ (std) | Avg # of genomes | Relative # of alignments |
|---|---|---|---|
| **Bacillales** | 9.98 | 38 | 100% |
| **Bacillales** | 9.59 ( 0.0651) | 11 | 7.17% |
| **Enterobacterales** | 24.92 | 145 | 100% |
| **Enterobacterales** | 28.33 (0.916) | 11 | 0.560% |
| **Methanobacteriales** | 7.46 | 11 | 100% |

**Table S1: Subsampling data impact on $\kappa$ estimation.** The subsampling is done so that Bacillales and Enterobacterales have the same average number of genomes per species than Methanobacteriales. The relative subsampling is the number of alignments necessary for the subsampled dataset divided by the original number of alignments. Inferred $\hat{\kappa}$ are robust to the amount of data used in the inference.

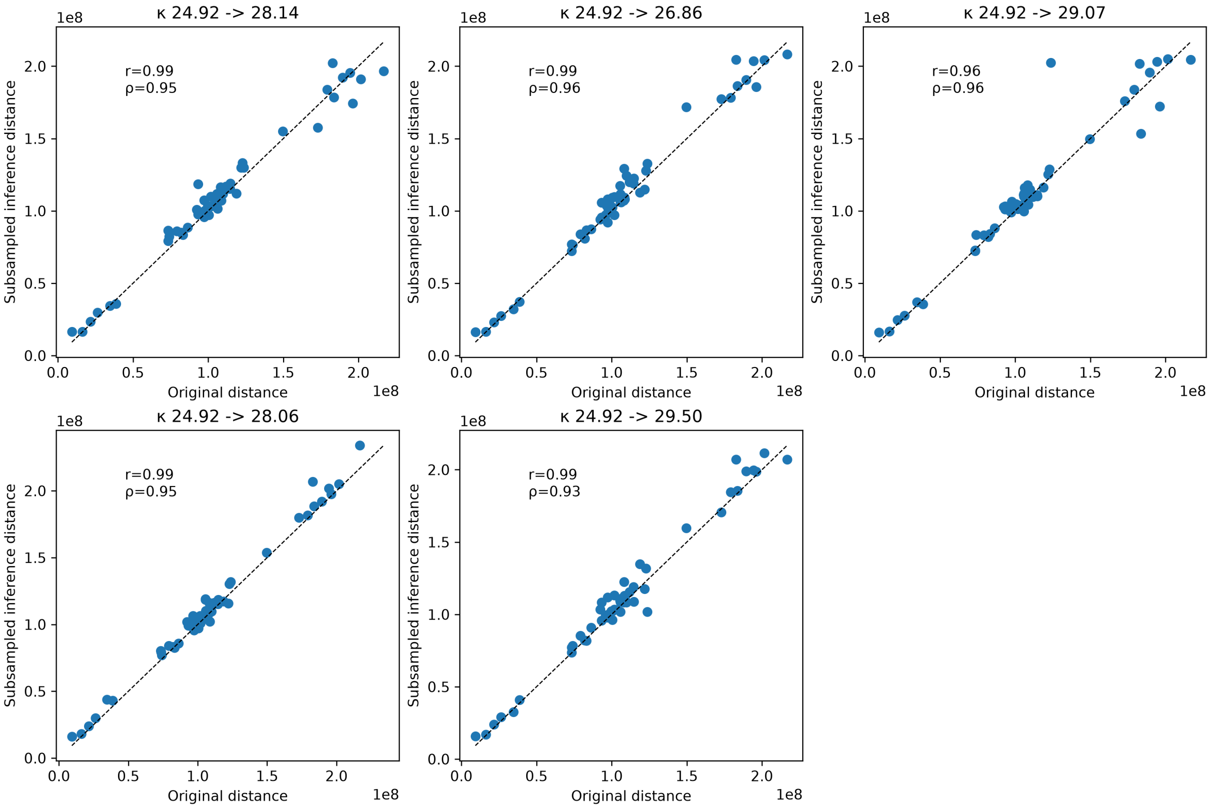


**Supplementary Figure S8: $\hat{\kappa}$ robustness to the amount of data.** *Enterobacterales* genomes are downsampled to the amount of data of Methanobacteriota. Each scatterplot shows the original distance inferred with full data and the distance inferred with subsampled data, along with Pearson and Spearman correlations. $\hat{\kappa}$ before and after subsampling are indicated in scatterplot titles. Each plot is one of five repeats to judge robustness.

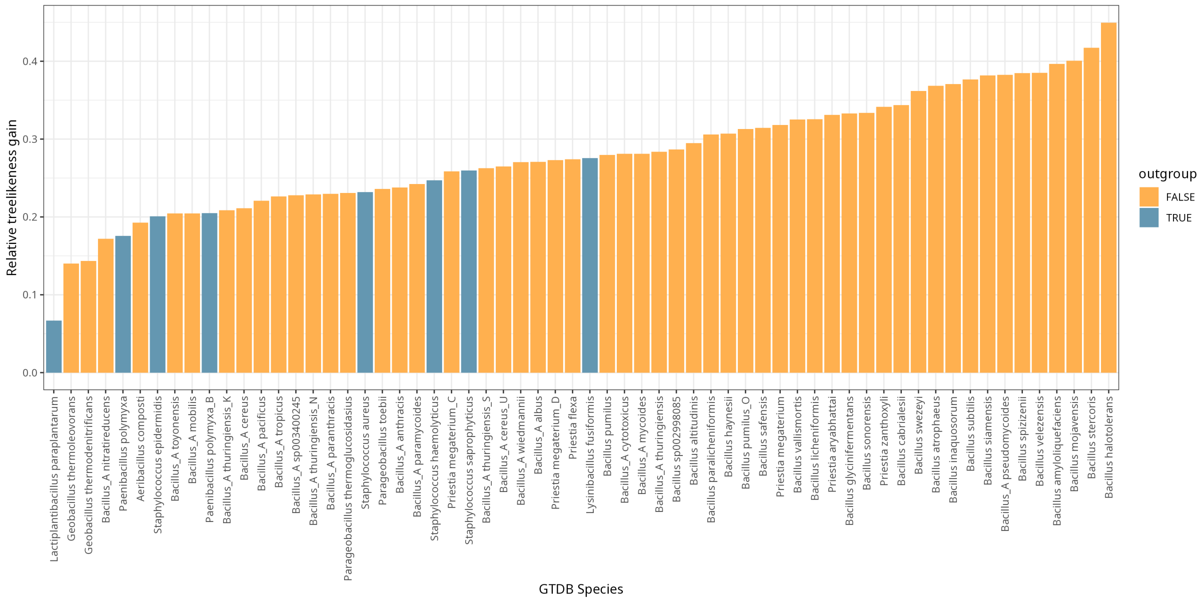


**Supplementary Figure S9: Gain of treelikeness for each *Bacillales* species from `mosaic-pair` to `mosaic-tree`.** The values are relative gain in average $\delta$ value across each quartet of leaves a given species participates in.

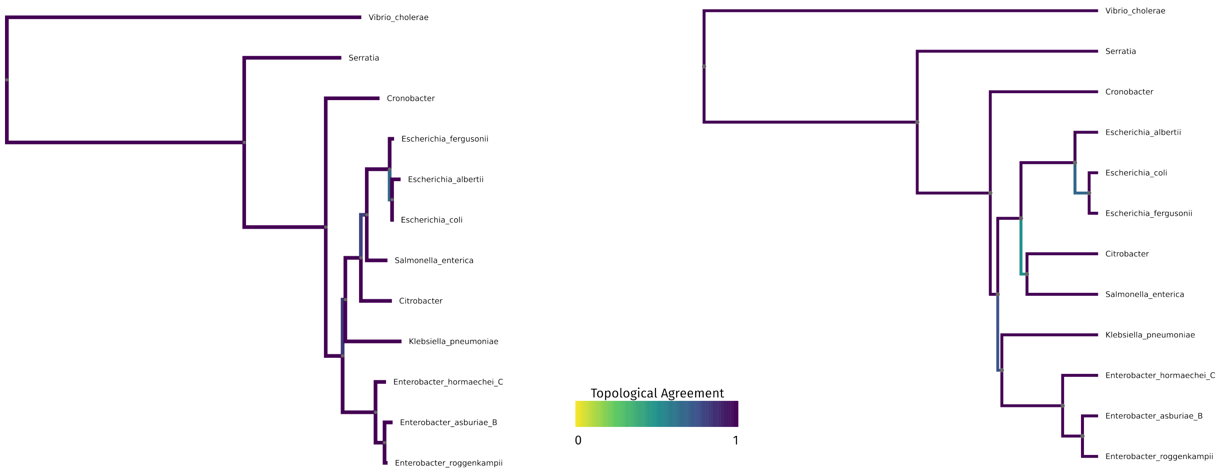


**Supplementary Figure S10: Reference GTDB tree (left) of *Enterobacterales*, inferred `mosaic-tree` tree (right).**

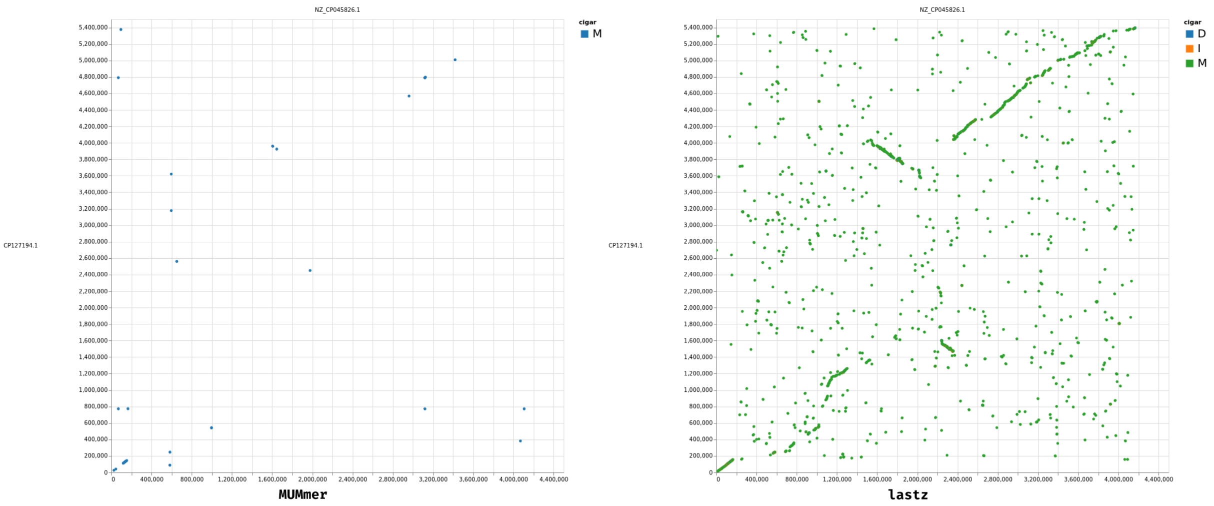


**Supplementary Figure S11: Dotplot of whole genome alignments by mummer (left) and lastz (right)** Pictured are a (*Bacillus_A bombysepticus* genome aligned to a *Bacillus subtilis* genome).

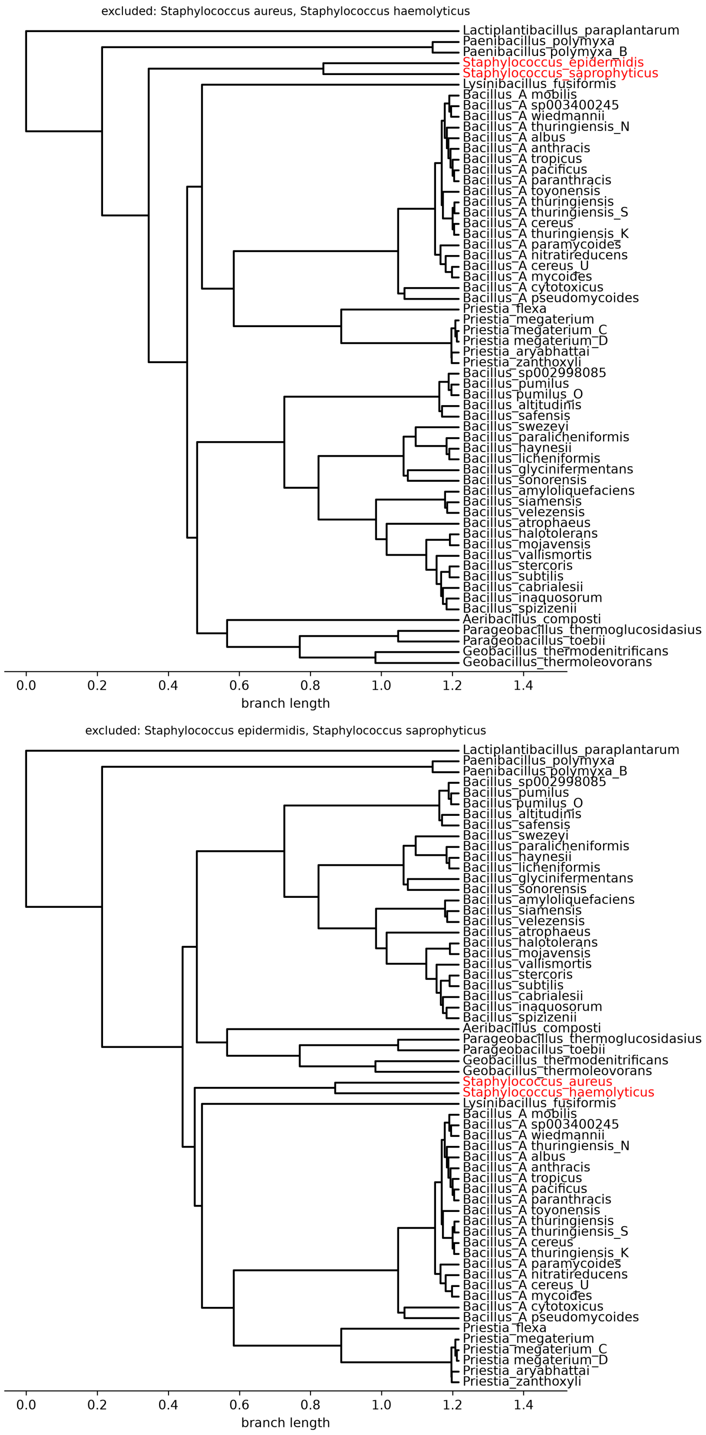


**Supplementary Figure S12: Two UPGMA trees excluding some *Staphyloccocales*.** UPGMA trees computed with `mosaic-tree` results on *Bacillales*, excluding the two close *Staphyloccocales* (*Staphylococcus aureus* and *Staphylococcus haemolyticus*) (top), and the two distant *Staphyloccocales* (*Staphylococcus aureus* and *Staphylococcus haemolyticus*) (bottom). The remaining *Staphyloccocales* are in red in both trees.

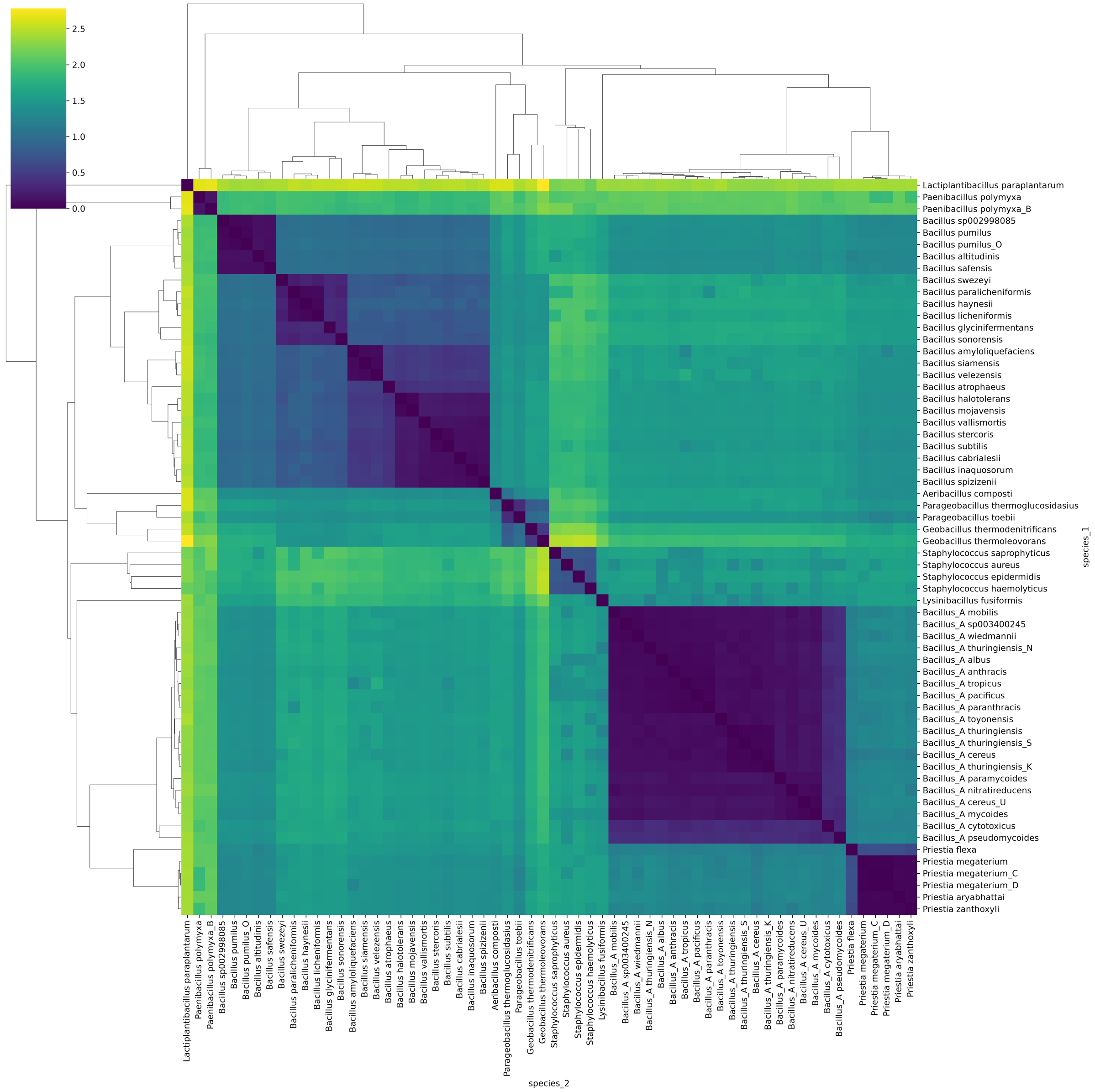


**Supplementary Figure S13: Heatmap and UPGMA trees of the *Bacillales* data**. Heatmap and UPGMA trees (row and column hierarchical clustering with average linkage) of the *Bacillales* data, representing the distance matrix inferred with `mosaic-tree`.

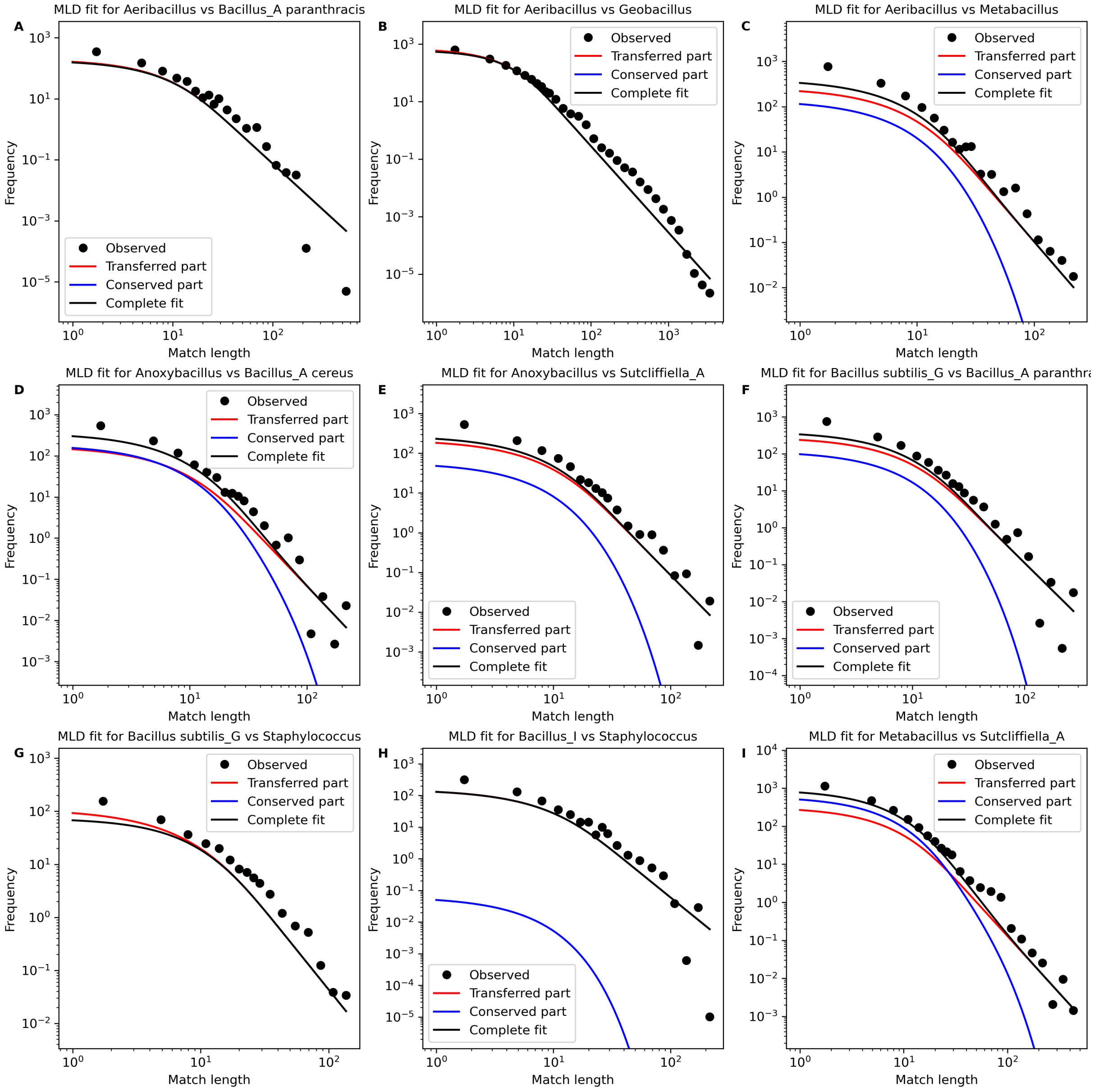


**Supplementary Figure S14: Examples of MLDs and fits from *Bacillaceae* MUMmer alignements**. Fits are separated in contributions of HGT (red) and clonal inheritance (blue). Some fits, for comparison of mostly distant species, do not contain a conserved component: A, B, E, F, G, H.

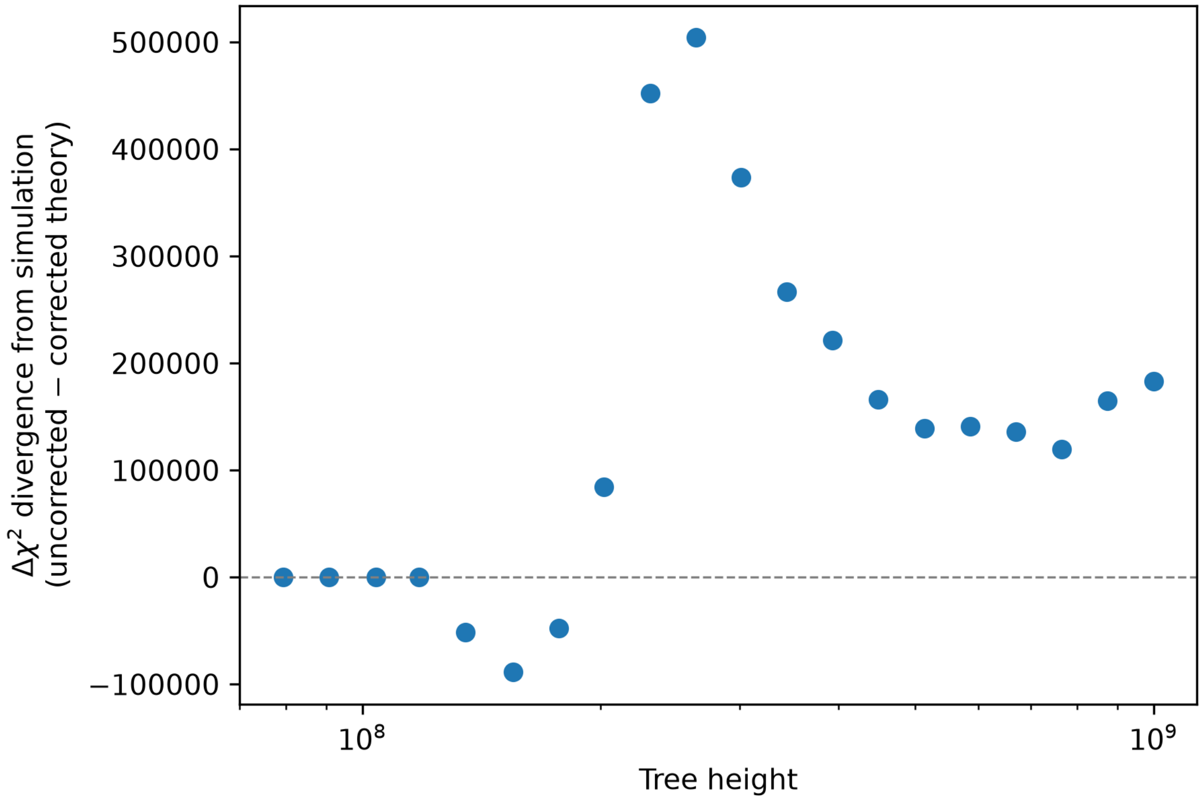


**Supplementary Figure S15: Improvement in model with the JC correction.** Difference in $\chi^2$ square divergence between the theoretical and simulated counts, in the uncorrected and corrected cases, for simulated pairs of genomes of increasing time divergence. Positive values indicate the JC-corrected theory is closer to the observed, simulated values. The JC69-corrected version of the aligner sensitivity fares better for longer elapsed time.
